# Gut bacteria translocation to the brain after ischaemic stroke occurs via the sympathetic nervous system

**DOI:** 10.1101/2023.04.03.535309

**Authors:** Alex Peh, Evany Dinakis, Michael Nakai, Rikeish R. Muralitharan, Samoda Rupasinghe, Jenny L. Wilson, Connie H.Y. Wong, Hamdi Jama, Charlotte M.O. Barker, Mahnaz Modarresi, Barbara K. Kemp-Harper, Tenghao Zheng, Francine Z. Marques, Brad R.S. Broughton

## Abstract

We provide evidence that stroke-induced gut breakdown results in bacteria translocation to the ischaemic mouse brain. Inhibition of sympathetic tone reduced bacterial load in the post-stroke brain and reduced functional deficits without altering cerebral apoptosis, neuroinflammation or infarct volume. These findings indicate that the activation of the sympathetic nervous system after stroke promotes gut-derived bacteria to enter to the brain, and this process worsens motor function in mice.

**Abstract Figure:** 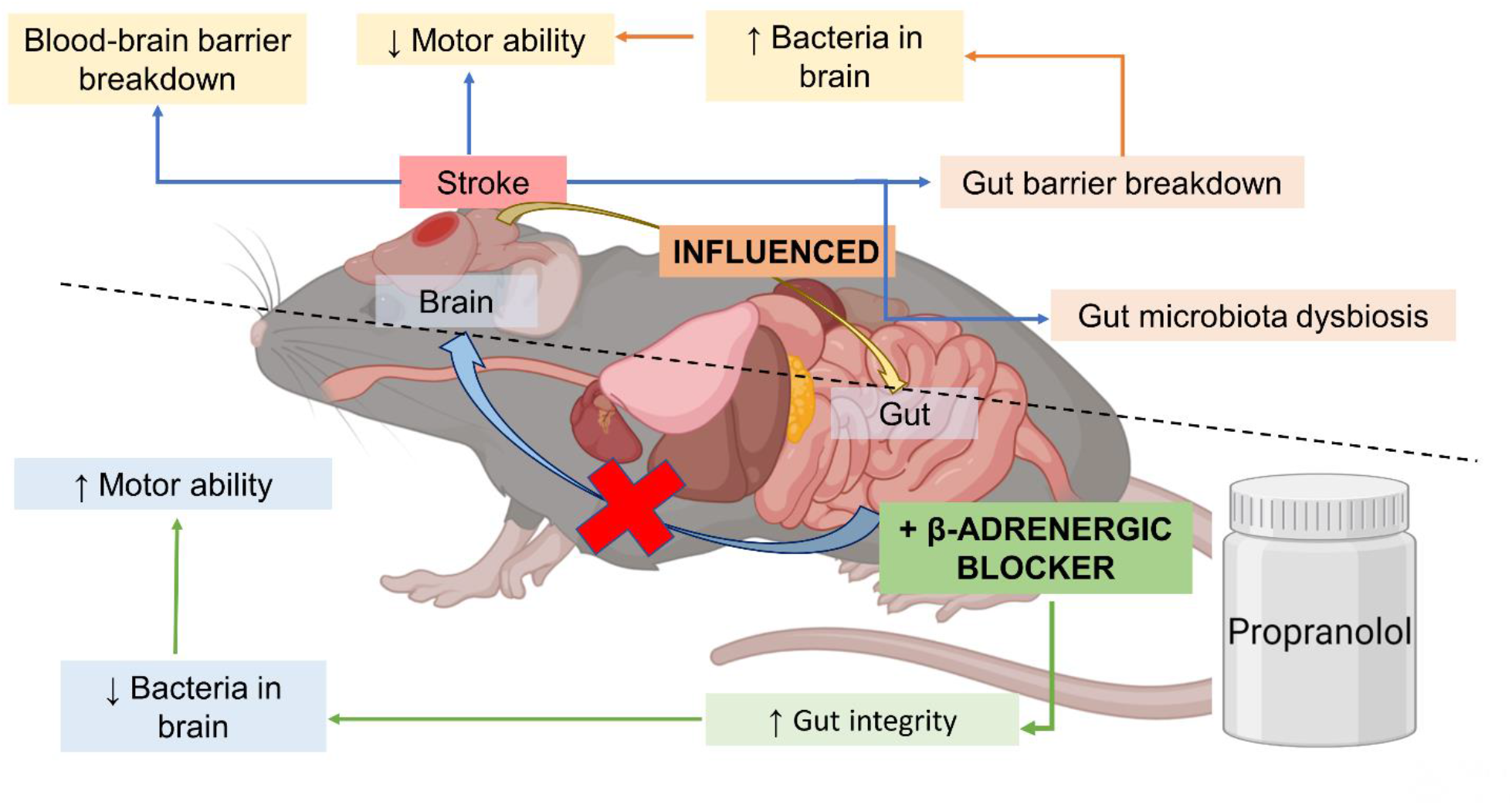

---

Infection is the most frequent complication of stroke, affecting approximately 30% of patients^1^, with pneumonia and urinary tract infections being the most common^2^. Bacterial infections after stroke are associated with neurological sequelae and prolonged hospital stays and are a significant cause of early- and long-term morbidity and mortality^3, 4^. Emerging evidence suggests that bacteria causing lung infection post-stroke may originate from the host’s gut^5^. Studies report that ischaemic stroke disrupts the gut-epithelial barrier and reduces gut bacterial diversity^5, 6^. Consequently, these changes likely contribute to the high incidence of poststroke infections, with translocation of bacteria from the gut to other systemic tissues, such as the lungs^6^. However, there is no evidence of whether gut bacteria migrate to the brain post-stroke. While the brain blood barrier (BBB) prevents bacteria from entering the brain under healthy conditions, it is severely compromised in the ischaemic hemisphere after stroke, thus increasing the likelihood of bacteria being able to enter the brain^7^.

We used two experimental mouse models of ischaemic stroke to determine whether bacteria infiltrates into the ischaemic hemisphere after 24 hours and provide a comprehensive characterisation of the acute intestinal changes post-stroke. Using Evans blue dye, we confirmed BBB breakdown in and around the infarct in our PT model (Fig. 1A,B). We observed a significant increase in peptidoglycan-immunoreactive bacteria in the ischaemic hemisphere of stroke mice (Fig. 1C,D), consistent with qPCR data that also detected increased bacteria in the ischaemic hemisphere (Fig. 1E) and blood of PT mice (Fig. 1F). Using 16S rRNA sequencing, we identified several genera of gut bacteria within the ischaemic brain post-PT stroke, including *Alistipes, Bacteroidales bacterium, Barnesiella_sp, Blautia, Lachnospiraceae NK4A136 group*, and *Parabacteroides* (Fig. 1G-L). We also observed significant peptidoglycan-positive bacteria in the brain of pMCAO specific free pathogen (SPF) but not in germ-free mice (Fig. 1M,N). To determine whether translocation of bacteria from the gut to the brain involves the sympathetic nervous system^6^, mice were treated pre- and post-stroke with the β-adrenergic receptor blocker, propranolol. While propranolol reduced bacteria translocation post-stroke (Fig 1C,D) and improved motor function (Fig. 2A), it did not affect infarct size, brain oedema, apoptosis, or neutrophil infiltration (Fig. 2B-H).

**Fig 1.**
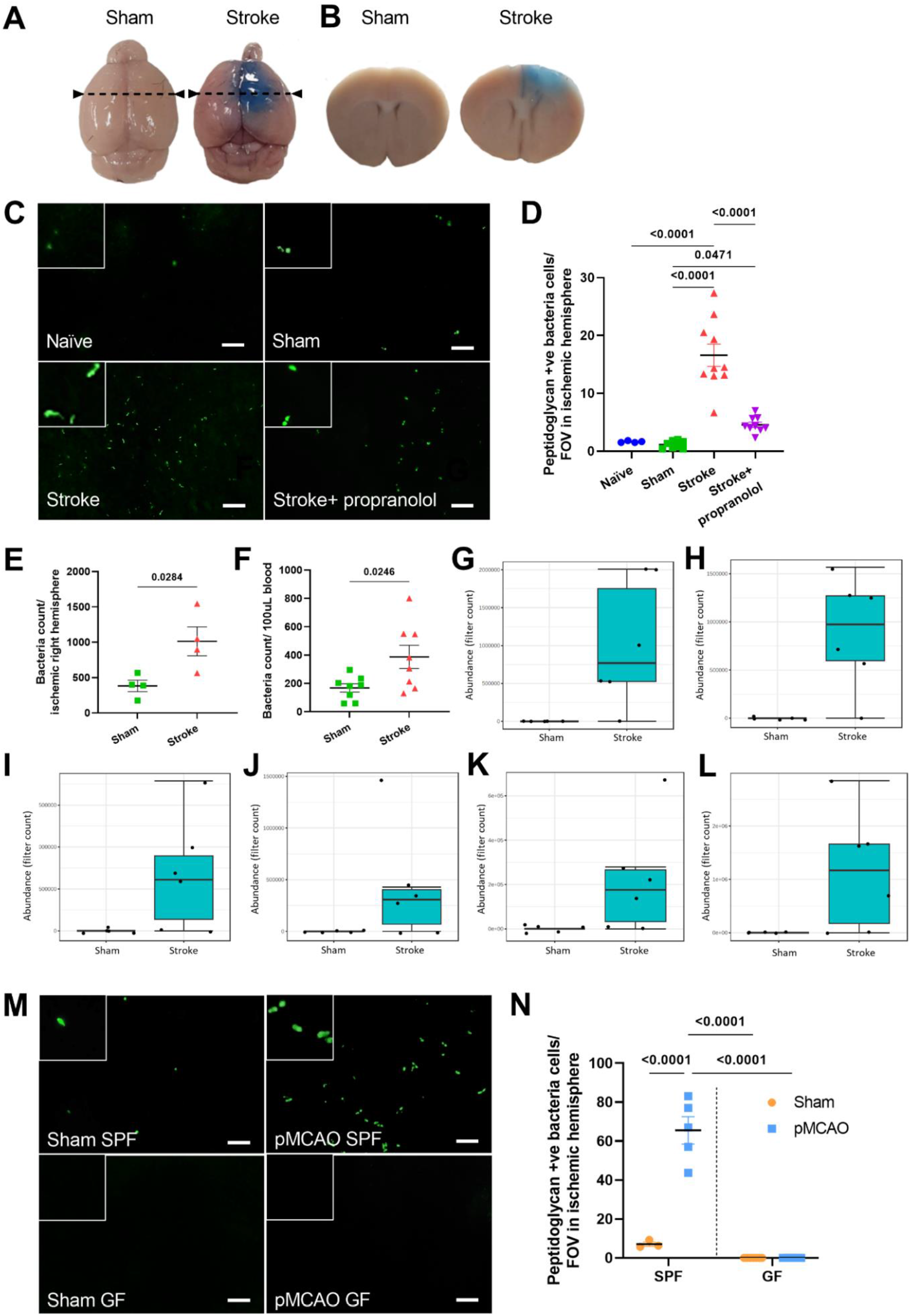
Detection of bacteria in the post-stroke brain as a result of BBB breakdown. Representative images of Evans blue dye extravasation in mice from a **(A)** superior view and **(B)** in corresponding coronal brain sections. Arrow lines indicate the location of coronal brain sections. **(C)** Representative image of positive peptidoglycan cells found in the ischaemic hemisphere of naïve, sham, PT stroke and propranolol-treated mice. Scale bar: 5µm. Inserts are magnified peptidoglycan-positive bacteria. **(D)** Quantification of peptidoglycan-positive cells in brain infarct region 24 hours after Photothrombotic (PT) stroke. Statistical test: One-way ANOVA corrected by false discovery rate (FDR); sample size=4-10/group. qPCR quantification of bacteria in **(E)** ischaemic right hemisphere brain and **(F)** blood of PT stroke mice. Statistical test: Student’s unpaired t-test; sample size= 4-10/group. Greater abundance of **(G)** *Parabacteroides* (P=0.090); **(H)** *Bacteroidales bacterium* (P=0.095); **(I)** *Barnesiella sp* (P=0.095); **(J)** *Blautia* (P=0.095); **(K)** *Alistipes* (P=0.095); **(L)** *Lachnospiraceae NK4A136 group* (P=0.095) were found in the brain of post-stroke mice using 16S microbiome sequencing method (n=5-6 mice/group). **(M)** Representative images of positive peptidoglycan cells were found in the pMCAO specific pathogen free (SPF), sham SPF, pMCAO germ-free (GF) and sham GF mice. Scale bar: 5µm. Inserts are magnified peptidoglycan-positive bacteria. **(N)** Quantification of peptidoglycan-positive cells in the brain infarct region 24-hours after permanent middle cerebral artery occlusion (pMCAO) stroke. Statistical test: Two-way ANOVA corrected by FDR. Sample size= 3-7/group. Error bars denotes mean±SEM.

**Fig 2.**
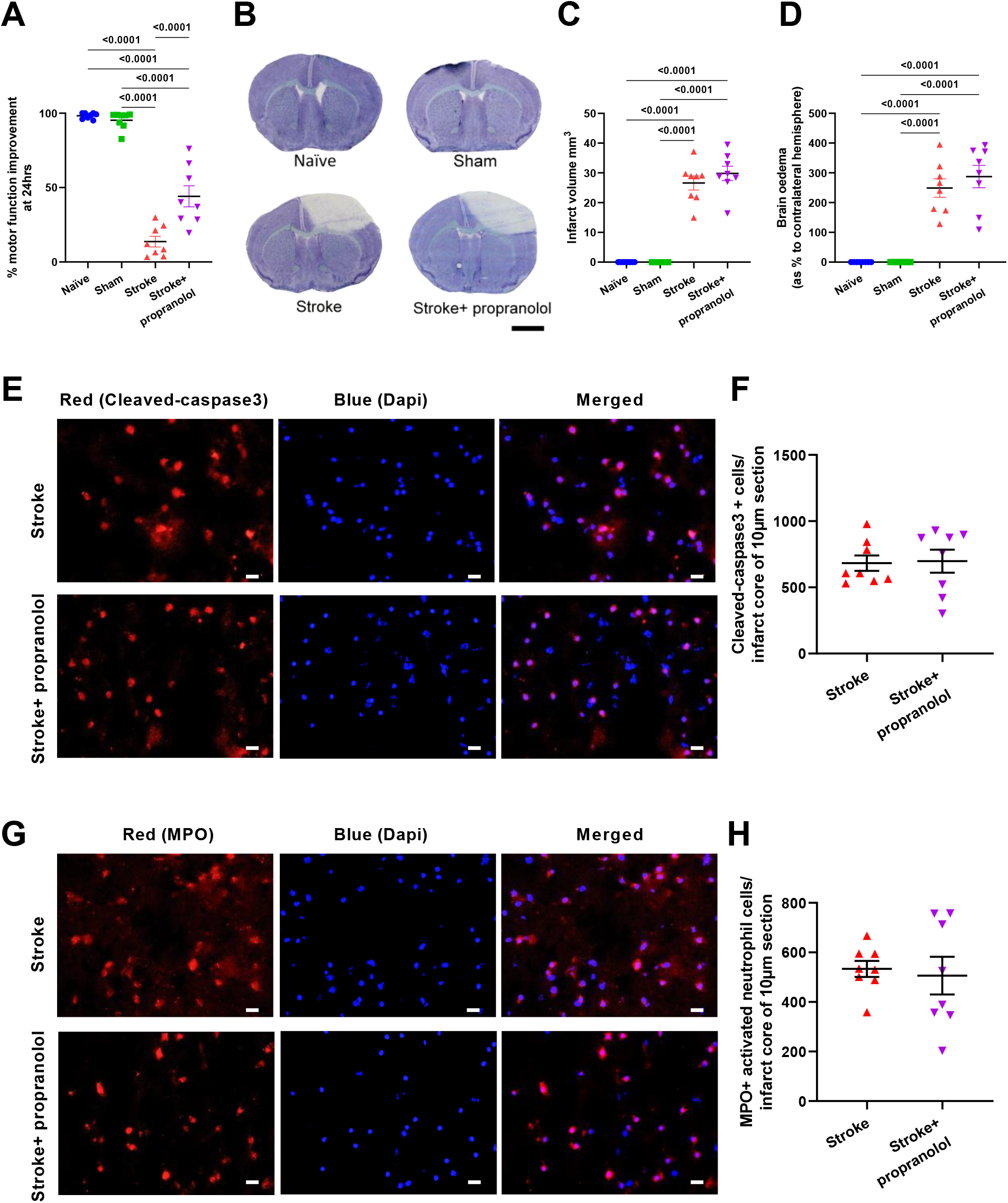
Role of the sympathetic nervous system (SNS) in bacteria translocation from the gut to the brain. **(A)** Data showing motor functional improvement assessed by wire-hanging test in four experimental groups. **(B)** Thionin-stained Photothrombotic (PT) stroke brain sections show the infarct region (white) and the unaffected area (purple). Scale bar: 2mm. **(C)** Infarct volume and **(C)** brain oedema of naïve, sham, stroke and propranolol-treated mice were measured at 24-hours poststroke. Statistical test: One-way ANOVA corrected by FDR. **(E)** Representative images of cleaved caspase-3 labelling in the infarct core of stroke and propranolol-treated mice. Scale bar = 20µm. **(F)** Quantification of cleavedcaspase-3 positive apoptotic cells in brain infarct region 24-hours after PT stroke using 10µm section. **(G)** Representative images of MPO-positive cells found in the infarct core of PT stroke and propranolol-treated mice. Scale bar = 20µm. **(H)** The number of MPO-positive cells in the infarct region quantified 24 hours post-stroke. Statistical test: Student’s unpaired t-test. Sample size= 8/group; error bar denotes mean±SEM.

Propranolol treatment also prevented gut barrier disruption following PT stroke. We found that there was a decrease in ZO-1 immunoreactivity in the colon following stroke, which was partially restored by propranolol treatment (Fig. 3A,B). Additionally, stroke increased mucus-producing goblet cells and reduced thickness of the muscularis propria within the small intestine (Fig. 3D-I) as well as shortening the length of the colon and small intestine (Fig. 3J-M). However, propranolol treatment consistently prevented these adverse effects (Fig. 3C-M). We also found increased gene expression related to gut barrier integrity and inflammation in the caecum (Supplementary Figure 1). However, no differences in villi length or fibrosis thickness were observed in the small intestine (Supplementary Figure 2) nor were differences in organ weights (Supplementary Figure 3).

**Fig 3.**
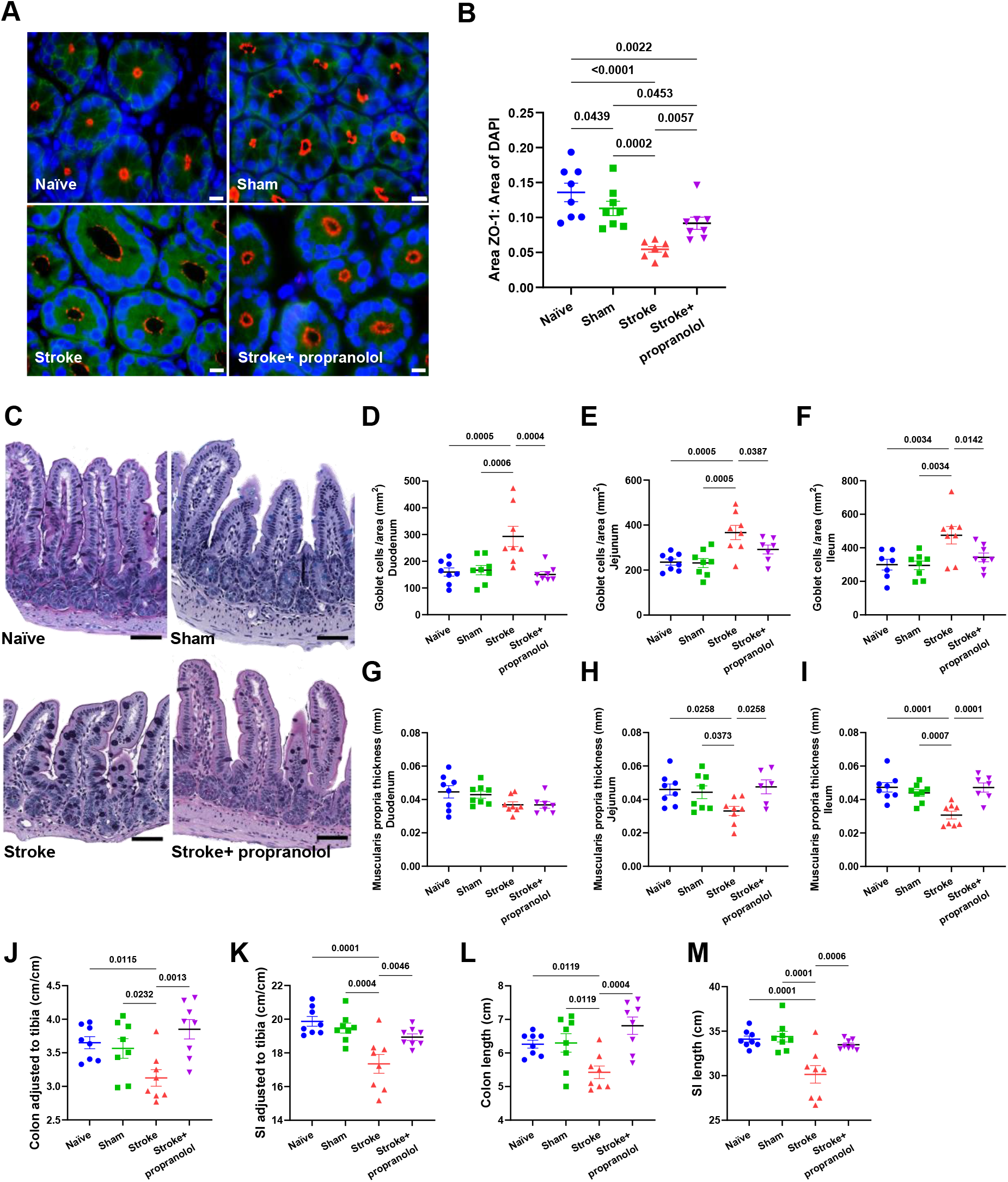
Stroke alters the gut-epithelial barrier. **(A)** Representative images of zonulin-1 (ZO-1) labelling in colon tissue of naïve, sham, stroke and propranolol-treated mice. Scale bar:10µm. **(B)** Quantification of ZO-1 expression on the colon tissue of naïve, sham, stroke and propranolol-treated mice. **(C)** Representative images the ileum depicting changes in the number of goblet cells and muscularis propria thickness stained using Periodic Acid-Schiff/Alcian blue. Scale bar: 60µm. Quantification of the number of goblet cells in the **(D)** duodenum, **(E)** jejunum and **(F)** ileum. Quantification of the thickness of muscularis propria in **(G)** duodenum, **(H)** jejunum and **(I)** ileum region. **(J)** The length of the colon and **(K)** the small intestine (SI) normalised to tibia length (cm/cm). **(L)** The length of the colon and **(M)** the SI measured 24-hours post-stroke. Statistical test: One-way ANOVA corrected with FDR. Sample size= 8/group; error bar denotes mean±SEM.

Furthermore, we analysed the changes in immune cells in various organs following PT stroke. We found increased immune cells in the brain (microglia, inflammatory monocytes, neutrophils) and blood (neutrophils) after stroke and decreased CD45+ immune cells in the mesenteric lymph nodes (Supplementary Figure 4). However, we did not find any changes in immune cells in the gut and spleen (see Supplementary Figure 5).

The impact of stroke on the gut microbiome was investigated using 16S rRNA bacterial gene analysis. PT mice were found to have less α-diversity in their gut microbiome than sham mice (Supplementary Figure 6A-D). The microbiome composition in the gut also differed between the two groups (Supplementary Figure 6E,F), with more Bacillota (formally Firmicutes) in sham mice and more Pseudomonadota (formally Proteobacteria) in stroke mice (Supplementary Figure 6G,H). Specific taxa that were more abundant in sham mice included the *Eubacterium xylanophilum group, Bacteroidales bacterium, Barnesiella_sp, Lachnoclostridium, Lachnospiraceae FCS020 group, Lachnospiraceae NK4A136 group, Lachnospiraceae_UCG_001, Lachnospiraceae_UCG_010, Roseburia, Ruminiclostridium*, and *Ruminococcaceae_UCG_014*. Conversely, *Enterococcus, Lactobacillus, Parabacteroides, Parasutterella*, and *Proteus* were more abundant in stroke mice (Supplementary Figure 6).

This study presents ground breaking evidence that gut bacteria infiltrate into the ischaemic hemisphere of two preclinical stroke models, highlighting the possibility of translocation from the gut to the brain via the breakdown of the gut-epithelial barrier and distribution through the host’s systemic blood circulation. The results also suggest that the β-blocker propranolol can prevent stroke-induced breakdown of the gut-epithelial barrier, thus reducing bacterial translocation to the brain, underscoring the significance of gut-sympathetic tone following ischaemic stroke.

Previous studies have reported the presence of bacteria in the brain of patients with neurological diseases, such as Alzheimer’s disease^8^. Additionally, research has identified bacteria in the blood clots of stroke patients, which are thought to come from the oral cavity^9^, but there is no evidence to suggest that these bacteria can cross the BBB. Thus, the current study is the first to demonstrate that the gut bacteria, including *Parabacteroides, Bacteroidales bacterium, Barnesiella sp, Blautia, Alistipes*, and *Lachnospiraceae NK4A136 group*, can migrate to the brain and cross the BBB post-stroke. It is noteworthy that while an increase in the levels of *Bacteroidales bacterium, Barnesiella sp*, and *Lachnospiraceae NK4A136 group* in the ischaemic brain was found, their levels were significantly lower in the gut after stroke when compared to the sham group, thus providing further evidence that the bacteria detected in the brain likely originated from the gut. Moreover, we observed no peptidoglycan-positive bacteria in the brain of germ-free mice post-MCAO, confirming that the bacteria originate from the gut.

The gut barrier relies on the mucus layer, muscularis layer, and intercellular tight junctions to maintain its integrity^10^. The current study found that cerebral ischaemia disrupts the gut barrier system, causing gastrointestinal inflammation that could contribute to the translocation of gut bacteria to the brain. Specifically, we observed thinner muscularis propria, increased goblet cell secretion, and decreased expression of the tight junction protein ZO-1 following PT stroke in mice. These findings are consistent with other animal models of ischaemic stroke^6, 11, 12^. However, genes responsible for maintaining gut integrity were upregulated, suggesting a compensatory response to restore gut barrier function. Moreover, an upregulation of various intestinal inflammatory genes was found post-stroke, indicating possible tissue damage.

It is well documented that an ischaemic stroke activates the sympathetic nervous and immune system, contributing to further brain injury and systemic complications^13^. We showed an increase in all major immune cell lineages within the ischaemic hemisphere post-PT stroke. It is possible that increases in microglia, resident and circulating monocytes and neutrophils in the brain post-stroke are in part due to bacterial infiltration, which may exacerbate post-stroke brain injury. The observed increase of immune cells in the ischaemic brain coupled with the decrease in the mesenteric lymph nodes following stroke suggests that the mesenteric lymph nodes may be the origin of some immune cells that migrate to the brain following a stroke, which is consistent with findings by Brea *et al*., (2021)^14^.

To investigate the impact of bacteria on brain damage following stroke, we used propranolol to block β-adrenergic signalling, which is known to cause the breakdown of the gut epithelial barrier^15^. We observed that propranolol treatment improved gut epithelial barrier integrity by increasing ZO-1 expression in the colon and muscularis propria thickness in the small intestine, as well as decreasing the number of goblet cell, thereby minimising gut bacterial translocation to the brain after stroke. Furthermore, propranolol treatment improved wire-hanging motor ability after stroke. Notably, we found that the reduction of bacteria in the brain was not associated with a decrease in neutrophil infiltration or apoptotic cells in the ischaemic hemisphere. Moreover, in infarct volume was not significantly reduced in mice treated with propranolol, which is consistent with a previous study comparing pMCAO germ-free mice and pMCAO SPF mice^6^. These findings indicate that the development and severity of the cerebral infarct volume after stroke are independent of bacteria within this region.

We acknowledge that this study has some limitations that should be considered. Our detection of bacteria in the brain provides only a static snapshot of bacteria present in that region at a specific time, as tissue analysis cannot capture the dynamics of bacterial translocation into the brain. While current whole-body imaging technology is available, its sensitivity is not optimal for detecting small numbers of translocated bacteria. In addition, we only examined the presence or absence of bacteria in the brain during the acute phase of stroke. Future studies should aim to track the location of the bacteria beyond this acute phase. Additionally, it is crucial to understand whether or not the presence of bacteria in the brain post-stroke is sex, age or strain dependent.

In conclusion, our study provides the first-ever evidence of bacteria in the brain post-stroke, likely to have migrated from the gut due to tight-junctions breakdown in the gut epithelial cells. While additional research is needed to determine whether the bacteria discovered in the brain are pathogenic or mutualistic, there is no indication that they play a role in causing acute brain damage.

## Acknowledgements

We would like to acknowledge the Monash Histology Facility for support with histology, the Monash FlowCore facility for support with flow cytometry, Monash Animal Research Platform for support with animal work, and the Monash Bioinformatics Platform for access to M3 servers. The Baker Heart & Diabetes Institute is supported in part by the Victorian Government’s Operational Infrastructure Support Program.

## Conflict of Interest

None.

## Funding

A.P is supported by Monash Graduate Scholarship and Monash International Tuition Scholarship. R.R.M is supported by a scholarship from the Faculty of Science, Monash University. S.R is supported by a Monash Graduate Scholarship. F.Z.M. is supported by a Senior Medical Research Fellowship from the Sylvia and Charles Viertel Charitable Foundation, a National Heart Foundation Future Leader Fellowship (105663), and National Health & Medical Research Council Emerging Leader Fellowship (GNT2017382). B.R.S.B is supported by a National Health & Medical Research Council Development Grant and a National Heart Foundation Grant.

## Author contributions

A.P. planned and performed most of the in vivo and in vitro animal experiments and data analyses, provided intellectual inputs and wrote the manuscript. E.D. contributed to in vivo animal works. M.N and H.J. contributed to 16S rRNA sequencing. R.R.M and S.R contributed to flow cytometry analysis. J.L.W and C.H.Y.W contributed to pMCAO surgery and GF mice study. B.K.K.H, F.Z.M., and B.R.S.B. contributed to study design. T.Z., F.Z.M., and B.R.S.B. conceived and supervised the research. F.Z.M., BKKH and B.R.S.B. secured funding to support this study. All authors approved the final version of the manuscript.

## Notes

### Competing Interest Statement

The authors have declared no competing interest.

