## Supplementary material for "Gut bacteria translocation to the brain after ischaemic stroke occurs via the sympathetic nervous system": Online supplemental tables and figures

**Short title:** Bacteria presence in the brain post-stroke

Alex Peh[1,2]; Evany Dinakis[1]; Michael Nakai[1]; Rikeish R. Muralitharan[1,3]; Samoda Rupasinghe[2]; Jenny L. Wilson[4]; Connie H. Y. Wong[4]; Hamdi Jama[1]; Charlotte M.O. Barker[2]; Mahnaz Modarresi[2]; Barbara K. Kemp-Harper[2]; Tenghao Zheng[1]; Francine Z. Marques[1,5*]; Brad R.S. Broughton[2*]

[1]Hypertension Research Laboratory, School of Biological Sciences, Monash University, Melbourne, Australia

[2]Cardiovascular & Pulmonary Pharmacology Group, Department of Pharmacology, Monash University, Melbourne, Australia

[3]Institute for Medical Research, Ministry of Health Malaysia, Kuala Lumpur, Malaysia

[4]Centre for Inflammatory Diseases, Department of Medicine, School of Clinical Sciences at Monash Health, Monash Medical Centre, Clayton, Victoria, Australia

[5]Heart Failure Research Group, Baker Heart and Diabetes Institute, Melbourne, Australia

*Contributed equally as senior authors

### Methods

#### Experimental studies

All the experimental procedures performed in this study were approved by the Monash Animal Ethics Committee (animal ethics approval number: MARP/2017/076, 23064, and MMCB/2018/002) and followed the Australian Code for the Care and Use of Animals for Scientific Purposes. Male C57BL6/J mice (8-9 weeks; n=4-12/group) were purchased from the Monash Animal Research Platform (MARP, Melbourne, Australia). Simple randomisation was used to group the mice into naïve, sham or stroke surgery. Researchers were blinded to the sample identification by using a unique sample ID for each animal. No mice were excluded from this study.

#### Ischaemic stroke model

The photothrombotic (PT) and permanent middle cerebral artery occlusion (pMCAO) stroke models were used in this study. PT stroke surgery was performed as previously described[1, 2]. Briefly, the mice were anesthetised with isoflurane, and Rose Bengal was injected into the peritoneum with a dose of 10mg/µl/g body weight. To induce photothrombosis, an area of 1.5mm diameter in the M1 cortex on the intact skull was focally illuminated for 15 minutes through a 25x objective to activate the dye. The mice were placed on a heating pad throughout the surgery, and body temperature was monitored via a rectal probe and maintained at 37°C. To confirm the presence of bacteria in the brain post-stroke, conventional and germ-free mice subjected to sham or pMCAO were used, as described in Shim *et al*.[3]. To induce pMCAO, a 10 mm incision was made on the right side of the neck after the mice were anaesthetised. A silicon-coated monofilament with a diameter of 0.21-0.23 mm was then inserted into the stump of the external carotid artery and ligated. The wound in the neck of the mice was sutured, and the mice were transferred to a heating pad to maintain their body temperature at 37 °C for recovery. To examine whether stroke-induced changes to the gastrointestinal (GI) tract occurs via sympathetic signalling mechanisms, PT mice were injected intraperitoneally with 250µg of propranolol (β adrenoreceptor blocker) 4 hours before and 6 hours after the PT stroke surgery.

**Wire-hanging test**

The wire-hanging test was performed as previously described[2], with slight modification to assess their motor function ability. In brief, the mice were hung on a wire stretched between two posts approximately 60cm high. The mice were able to utilise their forelimbs to hang their body weight on the wire. The time until each mouse dropped off the wire was recorded. Three hundred seconds was assigned as the maximum period, and three trials were performed for each mouse with 5 mins resting interval between trials. Their average hanging time was recorded. During the hanging wire test, if mice reached the posts, the timer was stopped, the animal was placed back in the middle of the wire and the timer was resumed.

**Sample collection**

All samples were collected following autoclaved-saline perfusion in a sterile environment (biosafety cabinet). All the equipment used for humanely killing the animals were autoclaved to remove any potential contamination. The mice were dipped in 70% ethanol after being euthanised. The head was decapitated, and the head was dipped in 70% ethanol again to ensure sterility. The large intestine and the spleen weight, as well as the small and large intestine length, were recorded. The duodenum, jejunum, and ileum tissue were collected, cut in half, and fixed in 10% neutral-buffered formalin. These tissues were paraffin-embedded and sectioned as well as stained for Periodic acid-Schiff/Alcian Blue (PAS/AB) and Masson's trichrome by the Monash Histology Platform. The tissues used for flow cytometry analysis were harvested and used immediately. In contrast, the caecal content, the brain, and the remaining parts of the gut were snap-frozen in liquid nitrogen and stored at -80$^{o}$C until needed for microbial DNA extraction or immunofluorescence, respectively.

**Infarct volume and brain oedema**

Frozen brains were cut into 30 mm coronal sections on a cryostat. Evenly spaced sections (separated by 210 µm for PT stroke or 420 µm for pMCAO) spanning the infarct were thaw-mounted onto poly-

L-lysine coated glass slides and stained with 0.1% thionin for infarct and brain oedema analysis. Brain oedema was calculated by comparing the ischaemic and contralateral hemispheres using the formula: Brain oedema (as % to contralateral hemisphere) = (volume of ischaemic hemisphere – volume of contralateral hemisphere)/ volume of contralateral hemisphere.

**Brain and colon immunofluorescence**

Serial coronal brain sections at 10 µm thickness were collected from the middle of the infarct for immunohistological analysis. All sections were thaw-mounted onto 0.1% poly-L-lysine coated glass slides and stored at -80°C. For colon sections, colon tissue was cut open along the mesenteric border, pinned flat without damaging the tissue and snap frozen in OCT (Tissue-Tek, Netherlands). All sections were then cryosectioned at 10um thick, thaw-mounted onto Superfrost-OT Plus glass slides (Thermo Fisher Scientific) and stored at -80°C. All immunofluorescence staining methods were performed after optimisation according to the manufacturer's protocol. Briefly, 10µm brain sections were air-dried before being fixed with 4% paraformaldehyde for 15 minutes. The sections were washed with 0.01M phosphate-buffered saline (PBS, 3x 10 minutes) and blocked with 10% goat serum (Abcam; Cambridge, USA, diluted in 0.01M PBS) for 1 hour. The sections were then incubated with a primary antibody overnight (see Supplementary Table 1), washed in 0.01M PBS (3x 10 minutes), and incubated with an appropriate secondary antibody at room temperature for 2 hours. The primary and secondary antibodies used are summarised in Supplementary Table 1. Sections were washed in 0.01M PBS (3x 10 minutes) and then carefully dried before being mounted in Vectashield Antifade Mounting Medium with DAPI (Vector Laboratories, Burlingame, CA). The same protocol was conducted for negative controls in the absence of primary antibodies and showed no immunofluorescence. To ensure that no cross-reactivity occurred with double-labelling immunofluorescence, controls consisted of all possible combinations of primary and secondary antibodies. A similar protocol was performed for the colon sections. Briefly, tissues were air-dried and fixed in 10% neutral-buffered formalin for 20 minutes, washed, labelled with primary antibody (see Supplementary Table 1) overnight, washed and labelled with secondary antibody (see Supplementary Table 1) for 2 hours at RT.

**Flow cytometry**

The brain, spleen, blood, mesenteric lymph nodes and gut were digested into single-cell suspensions by mechanical and/or enzymatic digestion with digestion buffer containing collagenase type XI (Sigma-Aldrich), hyaluronidase (Sigma-Aldrich), and collagenase type I-S (Sigma-Aldrich) as previously described[1]. To obtain a single cell suspension, samples were passed through a 70μm cell strainer, centrifuged and resuspended in fresh FACS buffer. We then stained the single-cell suspensions with various markers, as shown in Supplementary Table 2. The flow cytometry data acquisition was performed on the Fortessa X20 analyser (Becton and Dickson) at the Monash FlowCore (Australia). Data were analysed using FlowJo software version 10.

**qPCR quantification of bacteria in blood and brain**

The standard for the standard curve was made according to the manufacturer's protocol. In brief, bacterial 16S rRNA gene was used as the primer (F: 5'-CGGCAACGAGCGCAACCC; R: 5'-CCATTGTAGCACGTGTGTAGCC) to generate 16S PCR product. The PCR product was then cleaned, and the concentration was measured. The number of copies per ng DNA in PCR product was calculated using an online tool (http://www.thermoscientificbio.com/webtools/copynumber/). We then diluted the PCR product such that the solution contained $10^5$ copies of the gene per 1μL DNA. To generate the standard curve for quantifying bacteria in blood and brain, we performed a 10-fold serial dilution, starting with the $10^5$ sample. Real-time quantitative PCR (qRT-PCR) was performed in triplicates for each sample using Fast SYBR Green Master Mix (Thermo Fisher Scientific) on QuantStudio 7 real-time PCR system (Thermo Fisher Scientific) with one cycle of 95$^{o}$C for 10 min, followed by 40 cycles of 95$^{o}$C for 15 sec and 60$^{o}$C for 60sec.

**Microbial DNA extraction and 16S sequencing**

Both the ischaemic hemisphere of the brain and caecal content were collected sterilely during euthanasia and stored at -80$^{o}$C as previously described. To extract bacteria in the brain, the HostZero Microbial DNA Kit (ZymoResearch) was used following the manufacturer's protocol to first deplete the host DNA and then extract the microbial DNA. For the caecal content, we performed microbial DNA extraction using the DNeasy PowerSoil DNA isolation kit (Qiagen, Germany) according to the manufacturer's protocol. DNA from each sample was quantified using a Nanodrop (ThermoFisher Scientific). 10ng/μl of the microbial DNA was used to amplify the V4 region of the 16S ribosomal RNA, with 515F and 806R primers and Platinum™ Hot Start PCR Master Mix (ThermoFisher Scientific), according to the protocol of Earth Microbiome Project[4]. For brain samples, all DNA extracted was used as low numbers of bacteria were expected. 240ng of the library was pooled and purified using QIAquick PCR Purification Kit (Qiagen, Germany) before sequencing was performed at the Australian Genomic Research Facility (AGRF, Melbourne, Australia) on an Illumina MiSeq sequencer (300-bp paired-end reads).

**16S rRNA bioinformatics analyses**

The microbiome sequencing results were analysed using QIIME2 (2020.2 version) in R Studio (version 1.2.1335)[5]. The raw reads were first trimmed to have at least 20 Phred quality scores, merged the reads, and then removed the chimeric reads. DADA2 QIIME2 plug-in was used to remove and correct noisy reads[6]. Alpha-diversity metrics (ACE index, Chao1 richness index, Observed species richness, and Shannon diversity index), and beta-diversity metrics (both weighted UniFrac and unweighted UniFrac) shown as principle coordinate analysis (PCoA) plots were estimated using the q2-diversity QIIME2 plug-in after samples were rarefied for a minimum depth of 20,000 annotated sequence variants (ASVs). Data rarefying was not performed on brain samples due to the low count number. Taxonomy bar plots at phylum and genus levels, as well as Linear Discriminant Analysis (LDA) Effect Size (LEfSe) were visualised using MicrobiomeAnalyst[7].

**RNA isolation and qRT-PCR**

RNA from caecal tissue was extracted using TRIzol (Thermo Fisher Scientific), treated with DNase I (Thermo Fisher Scientific), and synthesised into cDNA using the High Capacity cDNA Reverse Transcription kit (Thermo Fisher Scientific) according to the manufacturer's protocol. qRT-PCR was performed in duplicates for each sample using Fast SYBR Green Master Mix (Thermo Fisher Scientific) on QuantStudio 7 real-time PCR system (Thermo Fisher Scientific) with one cycle of 95$^{o}$C for 20 sec, followed by 40 cycles of 95$^{o}$C for 1 sec and 60$^{o}$C for 20sec. Quantified genes and their primers are summarised in Supplementary Table 3. The qRT-PCR gene expression results were quantified using the $2^{(-\Delta\Delta Ct)}$ method[8]. Gene expression was expressed as fold change relative to caecal tissue from sham-operated animals. β-actin (*Actb*) was used as the reference gene.

**Statistical analyses**

Quantitative data were expressed as mean ± standard error of the mean (SEM). Statistical analyses were conducted using GraphPad Prism version 8 software (Graph-Pad Software Inc., San Diego, CA, USA). Statistical comparisons were made either by two-tailed unpaired t-test (when comparing two groups) or by one-way ANOVA corrected by FDR (when comparing more than two groups; two-stage step-up method of Benjamini, Krieger and Yekutieli) or by two-way ANOVA corrected by FDR (when comparing two grouping variables; two-stage step-up method of Benjamini, Krieger and Yekutieli). Results were expressed as mean ± standard error of the mean (SEM). $P<0.05$ was considered statistically significant.

#### Supplementary Tables

Supplementary Table 1. List of primary and secondary antibodies used for immunofluorescence.

| Tissue | Primary antibody | Dilution | Incubation temperature | Secondary antibody | Dilution |
|---|---|---|---|---|---|
| Brain | *Mouse anti-bacterial peptidoglycan (MAB995)* | 1:100 | Room temperature | Mouse on mouse Immunodetection Kit, Fluorescein (FMK-2201) | According to the manufacturer's protocol |
| | *Rabbit anti-MPO (ab9535)* | 1:200 | Room temperature | Goat anti-rabbit IgG Alexa Fluor 594 (Life Technologies) | 1:500 |
| | Rabbit Cleaved Caspase-3 (Asp175) (mAb9664) | 1:500 | 4˚C | Goat anti-rabbit IgG Alexa Fluor 594 (Life Technologies) | 1:500 |
| Colon (gut) | *Rabbit anti-mouse ZO-1 (61-7300)* | 1:100 | 4ºC | Goat anti-rabbit IgG Alexa Fluor 594 (Life Technologies) | 1:500 |
| | *Rat anti-mouse CD32/EpCam (14-5791-81)* | 1:500 | 4ºC | Goat anti-rat IgG Alexa Fluor 488 (Life Technologies) | 1:500 |

EpCam, epithelial cellular adhesion molecule; MPO, myeloperoxidase; ZO-1, zonula occludens-1

Supplementary Table 2. List of markers used for flow cytometry.

| Markers | Concentration | Conjugate | Fluorochrome | Clone | Manufacturer |
|---|---|---|---|---|---|
| Foxp3 | 1/500 | PE | YG585 | FJK-165 | eBio |
| CD4 | 1/500 | BV605 | BV610 | RM4-5 | Bioleg |
| CD8 | 1/500 | PerCP Cy5.5 | B710 | 53-6.7 | Bioleg |
| CD45 | 1/500 | AF700 | R730 | 30-F11 | Bioleg |
| Ly-6G | 1/1000 | PECy7 | YG780 | 1A8 | Bioleg |
| Ly6C | 1/500 | FITC | B530 | HK1.4 | Bioleg |
| F4/80 | 1/500 | APC-Cy7 | R780 | BM8 | Bioleg |

AF, Alexa Fluor; APC, Allophycocyanin; BV, Brilliant Violet; Cy, Cyanine; FITC, Fluorescein isothiocyanate; PE, Phycoerythrin; PerCP, Peridinin-Chlorophyll-Protein

Supplementary Table 3. A list of primer sets used for qRT-PCR.

| Pathway target | Gene name | Primer sequences |
|---|---|---|
| Gut epithelial integrity | Claudin 1 (Cldn1) | F: 5’- AAT TTC AGG TCT GGC GAC ATT |
| | | R: 5’- GGG GTC AAG GGG TCA TAG AA |
| | Mucin 2 (Muc2) | F: 5’- GCC CAC CTC ACA AGC AGT AT |
| | | R: 5’- GTC ATA GCC AGG GGC AAA CT |
| | Mucin 4 (Muc4) | F: 5’- TCC TCT TGC TAC CTG ATG CTC T |
| | | R: 5’- GCT CAT TTG GGA TGT TCT GGT G |
| | Tight junction protein 1 (Tjp1) | F: 5’- GGG CTC CTG GGT TTG GAT TT |
| | | R: 5’- GCA ACT CGG TCA TTT TCC TGT A |
| | Tight junction protein 2 (Tjp2) | F: 5’- AAA GCA GAG CCG AGC AAA TGG |
| | | R: 5’- GCT CTT GCG GAG GTT CTT CT |
| | Occludin (Ocln) | F: 5’- TTG AAC TGT GGA TTG GCA GC |
| | | R: 5’- AAG ATA AGC GAA CCT TGG CG |
| Inflammation | NLR family pyrin domain containing 3 (Nlrp3) | F: 5’- GGT GAC TTT GTA TAT GCG TGT TCT |
| | | R: 5’- GGG CTT AGG TCC ACA CAG AAA |
| | regenerating islet-derived protein 3β (Reg3β) | F: 5’- GGA AGA CAG ACA AGA TGC TGC |
| | | R: 5’- CTA ATG CGT GCG GAG GGT AT |
| | regenerating islet-derived protein 3ϒ (Reg3ϒ) | F: 5’- AGG ACA TCT TGT GTC TGT GCT |
| | | R: 5’- TCA TAG CCC AGT GTC GGG T |
| | Toll-like receptor 4 (Tlr4) | F: 5’- GGT AAG GTT GTC TTG ACG GAA C |
| | | R: 5’- GCC TCA GAG AAG GTA TCC AAC A |
| Housekeeping | β-actin (Actb) | F: 5’- AAC GGC TCC GGC ATG TGC AAA G |
| | | R: 5’-ATC ACA CCC TGG TGC CTA GGG CG |

#### Supplementary Figures

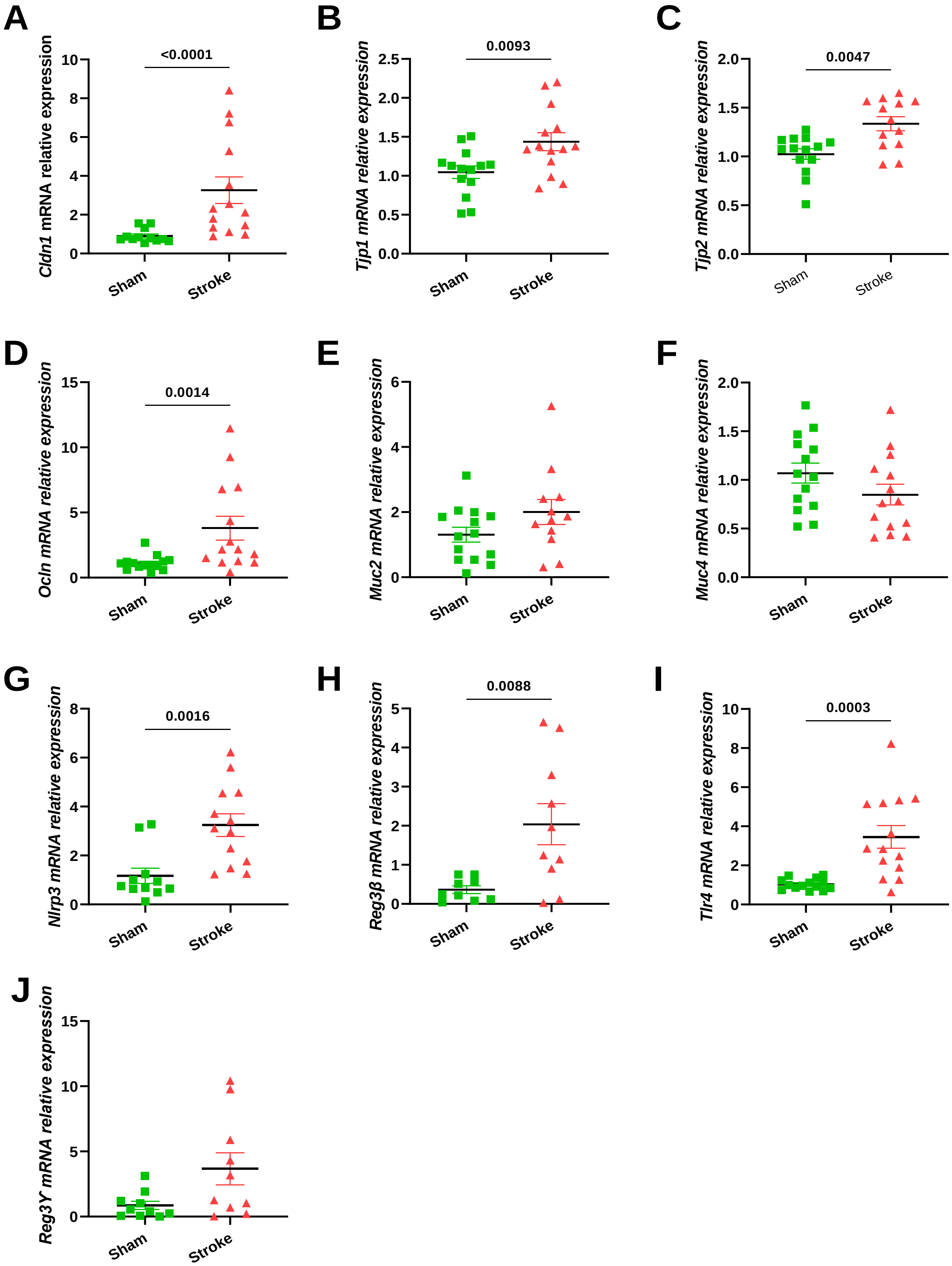


**Supplementary Figure 1. qPCR quantification of gene expression in caecal tissue.** The expression of genes in gut epithelial integrity: **(A)** *Cldn1*; **(B)** *Tjp1*; **(C)** *Tjp2*; **(D)** *Ocln;* **(E)** *Muc2*;

**(F)** *Muc4*, and inflammation: **(G)** *Nlrp3*, **(H)***Reg3β*, **(I)***Tlr4*, **(J)** *Reg3γ* of sham and PT stroke mice was performed. Statistical analysis: Two-tail unpaired-t test. Sample size= 9-14/group; error bar denotes mean±SEM.

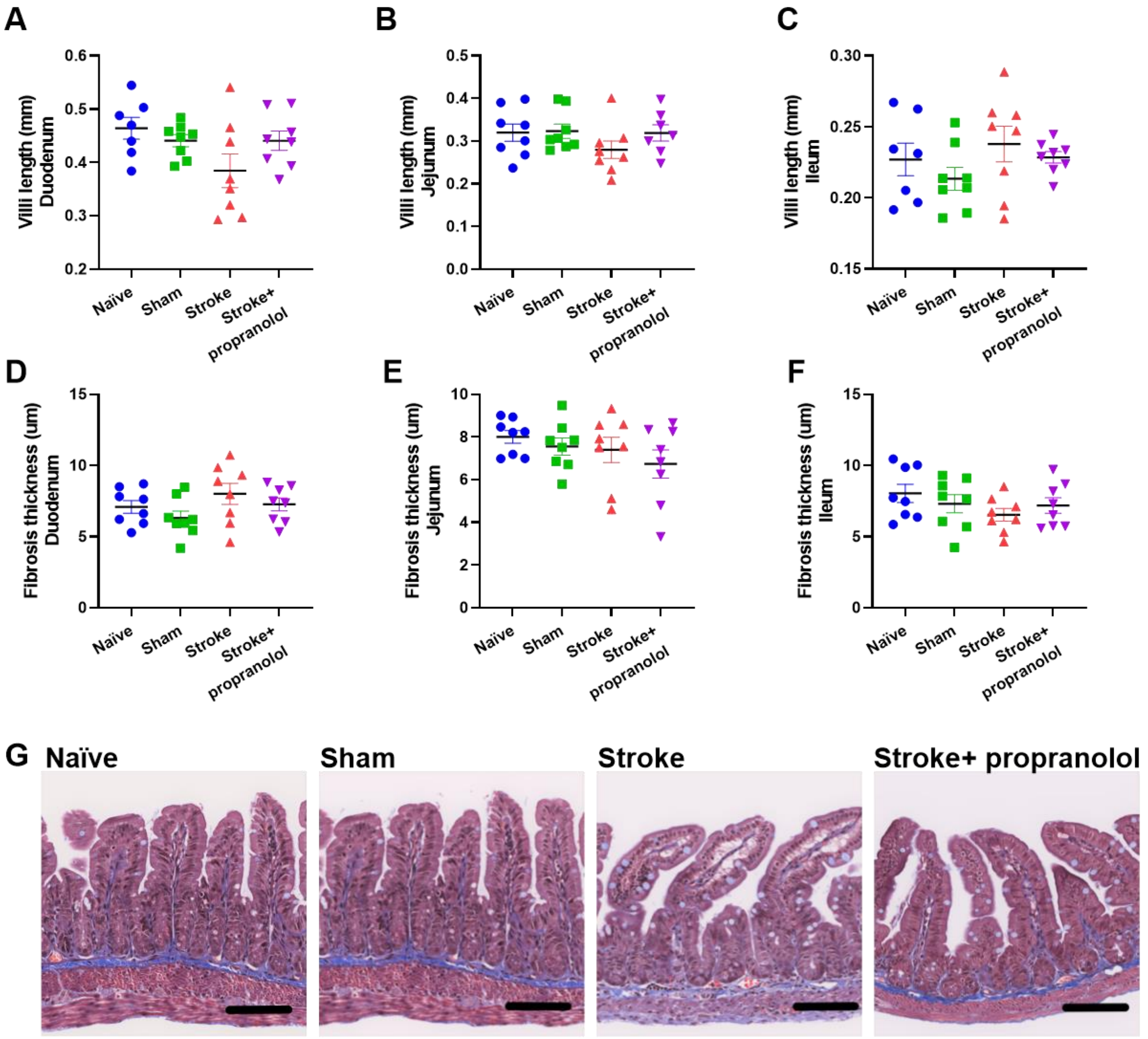


**Supplementary Figure 2. Villi length and fibrosis thickness between sham and stroke mice.** Quantification of the villi length in the **(A)** duodenum, **(B)** jejunum, and **(C)** ileum region, respectively. Quantification of the fibrosis thickness in the **(D)** duodenum, **(E)** jejunum and **(F)** ileum region, respectively. **(G)** Representative images showing the fibrosis thickness (region in blue colour) and villi length in naïve, sham, stroke and propranolol-treated mice. Scale bar: 100µm. Statistical test: One-way ANOVA corrected with FDR. Sample size= 8/group; error bar denotes mean±SEM.

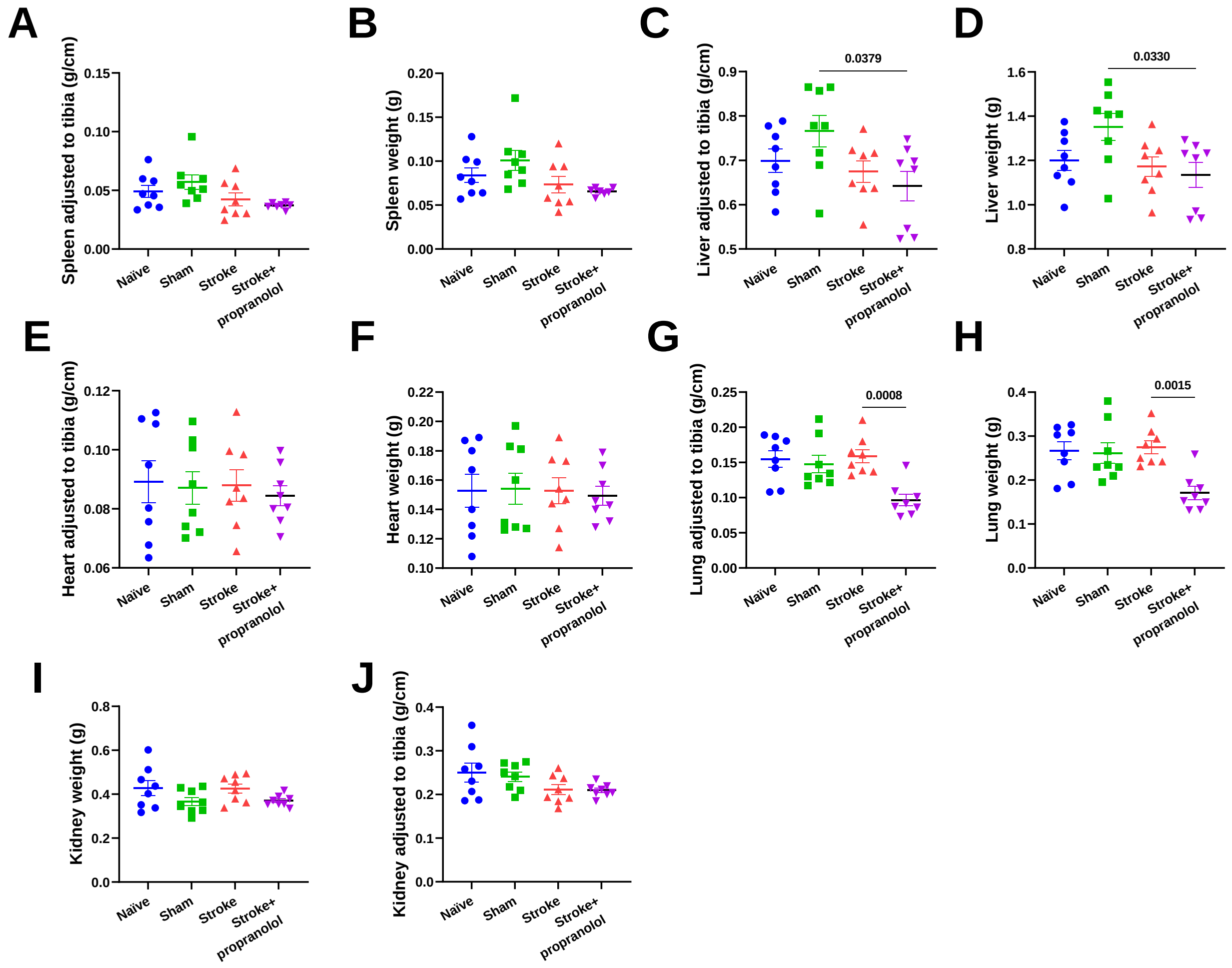


**Supplementary Figure 3. Organ weights between groups.** The raw weight of **(A)** spleen, **(B)** liver, **(C)** heart, **(D)** lung, and **(E)** kidney was measured 24-hours after surgery. The adjusted weights to tibia length of: **(F)** spleen, **(G)** liver, **(H)** heart, **(I)** lung, and **(J)** kidney was calculated. Statistical test: One-way ANOVA corrected with FDR. Sample size= 8/group; error bar denotes mean±SEM.

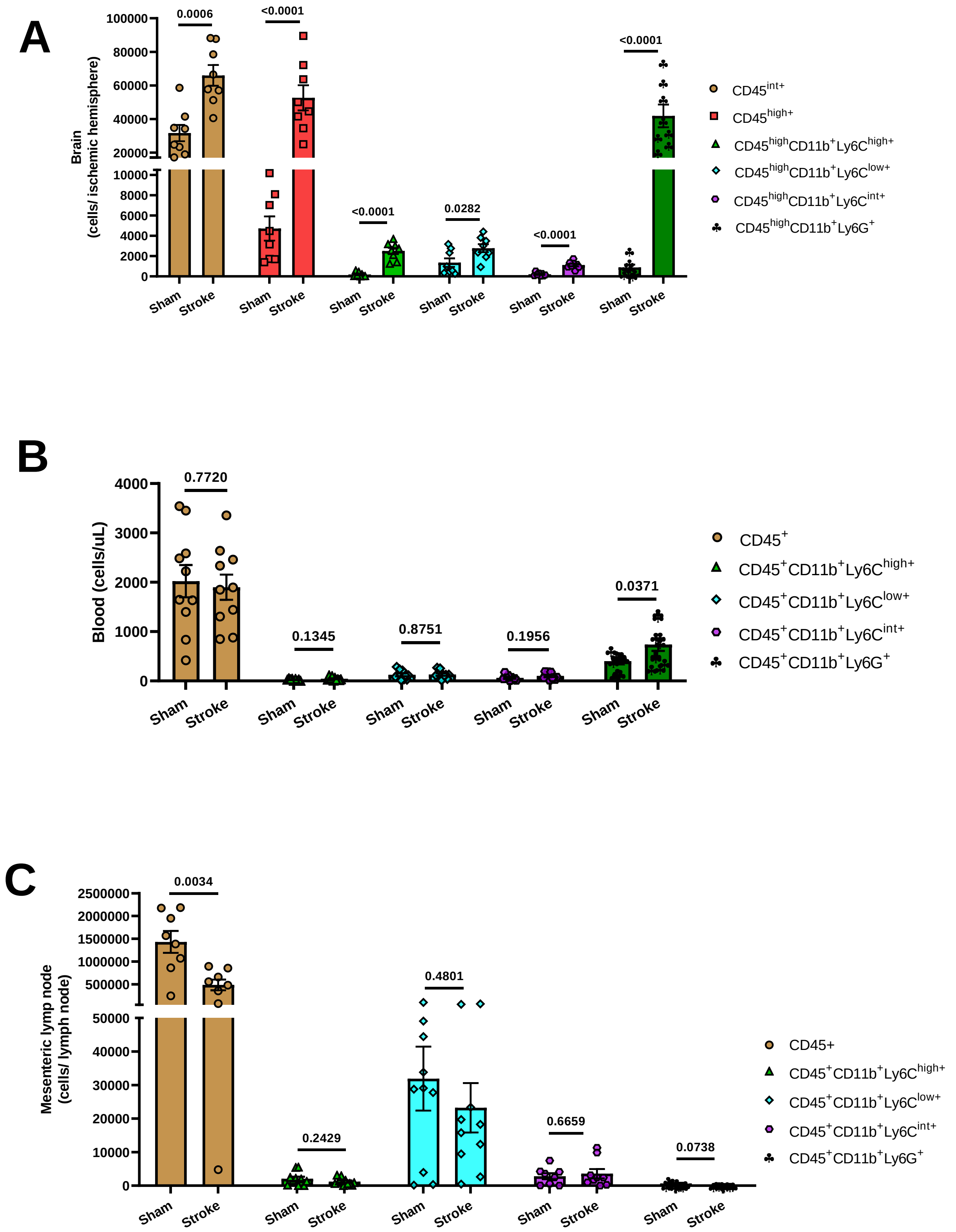


**Supplementary Figure 4. Changes in immune cell counts in various organs following stroke.**

Immune cell infiltration was assessed in **(A)** the brain; **(B)** blood; and **(C)** mesenteric lymph node between sham and stroke mice 24-hours following surgery using flow cytometry. Statistical test: Student’s unpaired t-test. Sample size= 8-14/group; error bar denotes mean±SEM.

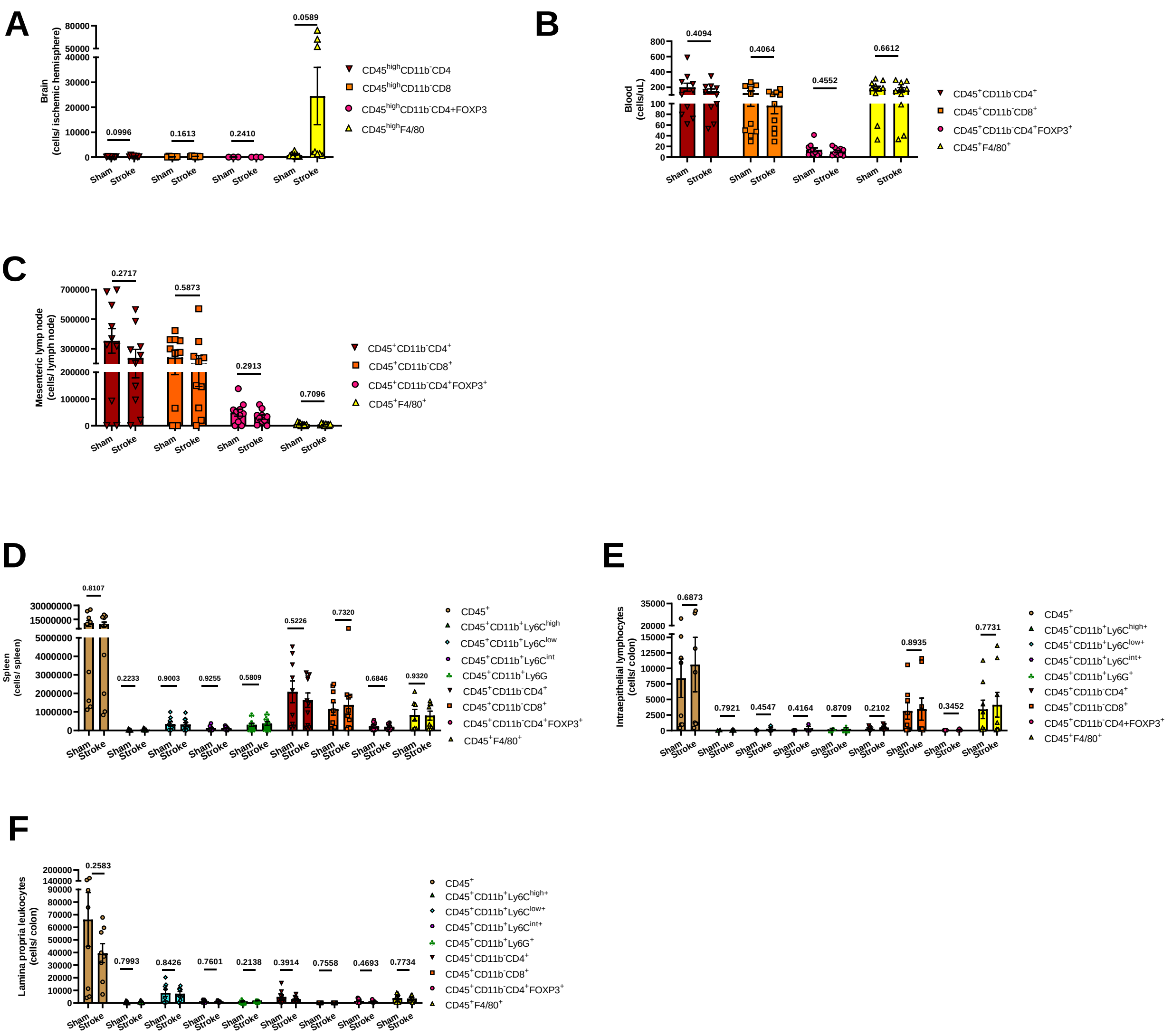


**Supplementary Figure 5. Changes in immune cell counts in various organs.** No significant change in immune cell counts between sham and stroke vehicle mice was found in **(A)** brain, **(B)** blood, **(C)** mesenteric lymph node, **(D)** spleen, **(E)** intraepithelial layer, and **(F)** lamina propria layer. Statistical test: Student’s unpaired t-test. Sample size= 3-10/group; error bar denotes mean±SEM.

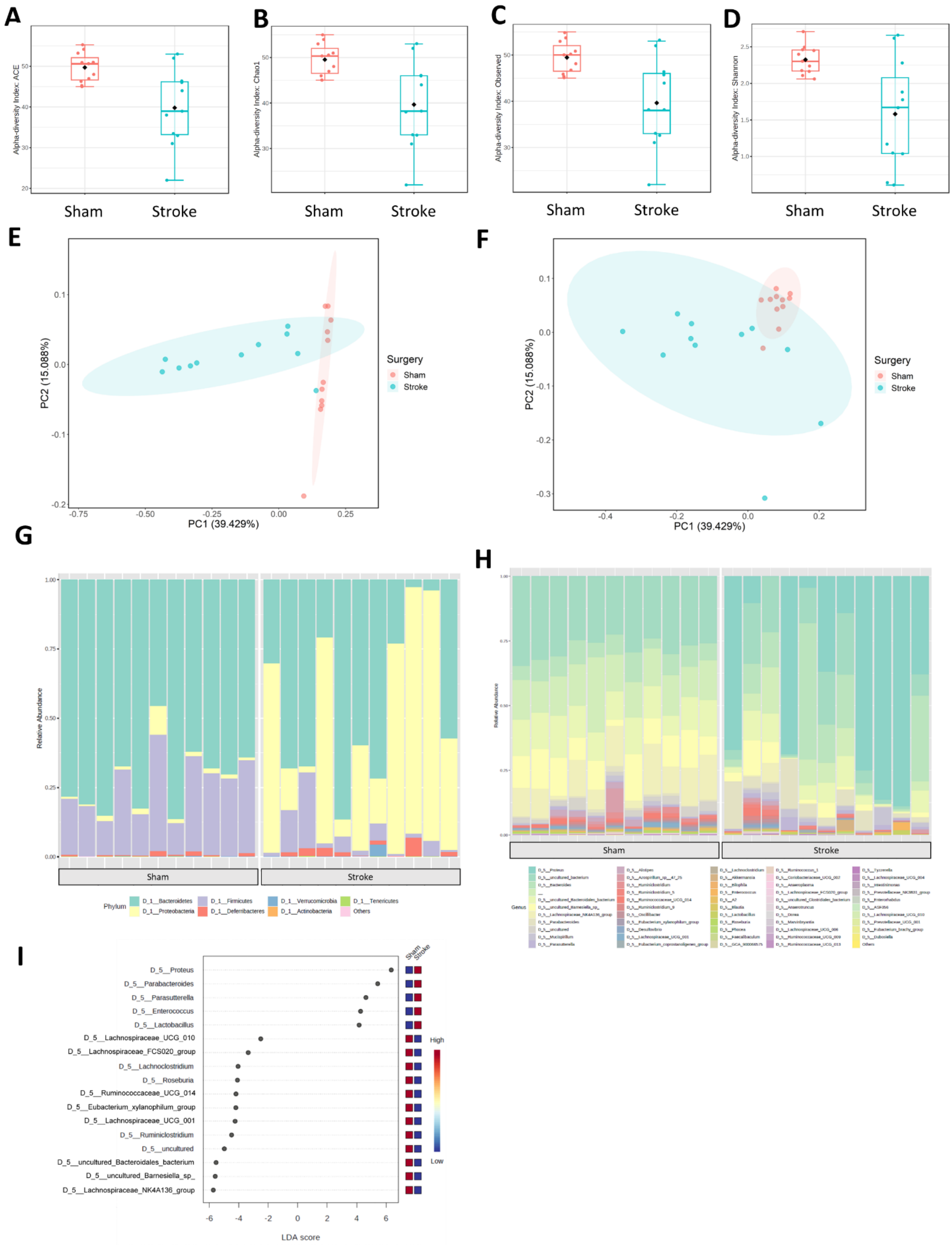


**Supplementary Figure 6. α- and β-diversity profiles of the gut microbiome.** Box plots illustrating 95%CI α-diversity of the gut microbiome between sham and stroke mice: **(A)** ACE index (p-value: 0.024); **(B)** Chao1 richness index (p-value: 0.020); **(C)** Observed species richness (p-value: 0.022); and **(D)** Shannon diversity index (p-value: 0.011). Principal coordinate analysis (PCoA) plots showing the clustering pattern of 95% CI β-diversity samples according to the type of surgery, based on **(E)** weighted UniFrac distance (P-value: 0.001); and **(F)** unweighted UniFrac distance (P-value: 0.003). Taxa bar plots classified based on **(G)** phylum- and **(H)** genus-levels. **(I)** Linear Discriminant Analysis (LDA) Effect Size (LEfSe) bar plot identified the top significant features of stroke vs sham (threshold on the logarithmic LDA score for discriminative features: 2.0; P-value cut off: 0.05, FDR-adjusted). Sample size= 11/group.
